# Standardization and validation of a panel of cross-species microsatellites to individually identify the Asiatic wild dog (*Cuon alpinus*): implications in population estimation and dynamics

**DOI:** 10.1101/447201

**Authors:** Shrushti Modi, Bilal Habib, Pallavi Ghaskadbi, Parag Nigam, Samrat Mondol

## Abstract

**Background:** The Asiatic wild dog or dhole (*Cuon alpinus*) is a highly elusive, monophyletic, forest dwelling, social canid distributed across south and Southeast Asia. Severe pressures from habitat loss, prey depletion, disease, human persecution and interspecific competition resulted in global population decline in dholes. Despite a declining population trend, detailed information on population size, ecology, demography and genetics is lacking. Generating reliable information and landscape level for dholes is challenging due to their secretive behaviour and monomorphic physical features. Recent advances in non-invasive DNA-based tools can be used to monitor populations and individuals across large landscapes. In this paper, we describe standardization and validation of faecal DNA-based methods for individual identification of dholes. We tested this method on field-collected dhole faeces in four tiger reserves of the central Indian landscape in the state of Maharashtra, India. Further, we conducted preliminary analyses of dhole population structure and demography in the study area.

**Results:** We tested a total of 18 cross-species markers and developed a panel of 12 markers for unambiguous individual identification of dholes. This marker panel identified 101 unique individuals from faecal samples collected across our pilot field study area. These loci showed varied level of amplification success (57-88%), polymorphism (3-9 alleles), heterozygosity (0.23-0.63) and produced a cumulative probability of identity _(unbiased)_ and probability of identity _(sibs)_ value of 4.7×10^−10^ and 1.5×10^−4^, respectively. Our preliminary analyses of population structure indicated four genetic subpopulations in dholes. Qualitative analyses of population demography show signal of population decline.

**Conclusion:** Our results demonstrated that the selected panel of 12 microsatellite loci can conclusively identify dholes from poor quality, non-invasive biological samples and help in exploring various population parameters. Our methods can be used to estimate dhole populations and assess population trends for this elusive, social carnivore.

## Introduction

The Asiatic wild dog or dhole (*Cuon alpinus*) is a highly elusive, endangered, social canid distributed in south and southeast Asia [1, 2] occupying a range of habitat types including alpine, temperate, sub-tropical and tropical forests [2]. Driven by habitat loss, prey depletion, disease transmission from domestic dog, human persecution and interspecific competition [3, 4], dholes are currently found in about 75% of their historical global range [2, 4]. Global dhole population is roughly estimated to be about 4500-10500 with only 949-2215 mature individuals, but accurate estimates and population trends are not available from any part of its range [4]. They are considered as ‘Endangered’ by IUCN under criteria C2a(i) and Appendix II of the Convention on International Trade in Endangered Species (CITES). The Indian subcontinent currently retains majority of the remaining dhole populations [4], where the species has faced about 60% decline in their historical distribution [5]. The Western Ghats and central Indian forests of the subcontinent still holds majority of the remaining dhole population in India[6], whereas small populations are found in the Eastern Ghats [6], northeast India [7, 8] and Himalayan region [9]. Given the current anthropogenic disturbance scenario across its range, the future survival of this monotypic genus depends on integrated conservation measures involving detailed, accurate information on ecology, demography and genetics.

However, generating reliable information for this elusive, forest-dwelling and pack-living canid at landscape scale is challenging. Traditional ecological techniques such as regular photographic capture approach is ineffective for dholes due to absence of unique coat patterns and their monomorphic forms, and physical tagging methods are impractical at landscape scales due to logistical difficulties, high costs and small numbers of captures possible. In this context, genetic tools have tremendous potential to generate critical information (for example, population size estimation [10], phylogeography [11, 12], pack dynamics and reproductive fitness [13, 14], dispersal patterns [15, 16] etc.) for elusive species conservation across large landscapes [17]. The ability to identify individuals from non-invasive samples collected over large space provide a feasible option to generate detailed information on elusive, forest-dwelling dholes as they cannot be identified using other approaches.

In this study, we addressed key methodological issues related to selection and standardization of a set of molecular markers for individual identification of dholes. Subsequently, we tested these markers on field-collected dhole samples from three tiger reserves of the central Indian landscape in the state of Maharashtra, India for individual identification and conducted preliminary analyses of population parameters this this landscape. In addition to the utility of these markers in dhole population estimation at landscape level, we believe that this approach has wider relevance in non-invasive, fecal DNA based population assessments of many other low density, elusive, wide-ranging species.

## Methods

### Research permits and ethical considerations

All required permissions for fieldwork and sampling were provided by the Maharashtra Forest Department (Permit No. 09/2016). The entire study was non-invasive through field-collected faecal samples, and thus did not require any ethical clearance from the institute.

### Study Area

The study was focused in five protected areas Melghat Tiger Reserve (MTR), Pench Tiger Reserve (PTR), Navegaon-Nagzira Tiger Reserve (NNTR), Tadoba-Andhari Tiger Reserve (TATR) and Umred-Karandhla Wildlife Sanctuary (UKWLS) of the central Indian landscape in the state of Maharashtra, India. The entire area is a complex of forested areas (core zone) with different levels of connectivity. NNTR and PTR are geographically closer as compared to MTR-NNTR and MTR-PTR. MTR and PTR are part of the Satpura-Maikal-Pench corridor in the Satpura-Maikal landscape. The forest type is of dry deciduous to moist deciduous nature with major vegetation consisting *Tectona grandis, Anogeissus latifolia, Lagerstroemia parviflora, Terminalia spp., Heteropogon contortus, Themeda quadrivalvis, Cynodon dactylon* etc.

### Field Sampling

Dholes prefer dense forested habitats [18] where the social groups defecate in communal latrine sites [1]. Their elusive nature and highly social behaviour presents unique challenges in scat sampling for individual identification. In this study, sampling was conducted through intensive foot and vehicle surveys covering the entire study area to identify dhole latrine sites. Once the site is found, one bolus from each fresh scat was collected considering from one individual. Separate gloves were used to collect each sample. All samples were collected directly in wax paper and stored in separate ziplock bags. Once brought to the field station, the sample containing ziplock bags were temporarily stored in a large box containing silica gel to minimize fungal growth in humid conditions. Samples were then shipped to the laboratory, where they were stored in a −20°C freezer. GPS co-ordinates and other associated information (track marks, substrata etc.) were collected for each sample. Entire sampling was conducted between January 2015 – June 2017, covering PTR (257.3 km^2^), MTR (1500.49 km^2^), NNTR (152.8 km^2^), TATR (627.5 km^2^) and UKWLS (189 km^2^), Maharashtra. A total of 249 samples were collected (PTR- 92, MTR- 76, NNTR- 37, UKWLS- 34, TATR- 10, respectively) for this study. Details of all sample locations are given in Supplementary Figure 1.

In addition, four dhole blood samples were also collected as reference samples from Tadoba-Andhari Tiger Reserve in 2017-18 for standardisation of initial marker selection and individual identification. Blood samples were collected in EDTA vacutainers and subsequently stored in a −20°C freezer in the laboratory.

### DNA extraction

DNA extraction was performed in the laboratory from the frozen faecal samples using QIAamp DNA Tissue Kit (QIAGEN Inc., Hilden, Germany) with a modified approach, depending on sample quality. If the sample had the entire top mucous layer available (i.e. not covered by dust, soil etc.) then it was swabbed with phosphate buffer saline soaked sterile cotton swab and was stored in sterile Eppendorf tube at −20°C. However, if the mucous layer was covered then the top layer was scraped using sterile blades and stored in similar conditions. Subsequently, faecal samples collected by both methods were lysed overnight in 300/600 μl of lysis buffer for swabs and scraped samples, respectively and 20 μl proteinase K followed by extraction using the kit’s protocol. DNA was eluted twice with 100 μl of 1X TE and stored in −20°C for long-term use. Each set of 22 extractions was accompanied with two negative controls to monitor possible contamination.

DNA from blood samples was extracted using standard protocol given in the QIAamp DNA Tissue Kit (QIAGEN Inc., Hilden, Germany). Negative control was incorporated to monitor any possible contaminations.

### Selection of microsatellite markers

There are no dhole specific microsatellites developed so far and the only study focusing on dhole population genetics had used 13 cross-species markers from domestic dogs to study genetic variation [19]. These markers showed low levels of polymorphism and low PID_sibs_ value (3.3×10^−4^), indicating a potentially inefficient panel for unambiguous individual identification at landscape levels with large population sizes. For this study, we developed a panel following stringent cross-species marker selection and testing process. The entire process was conducted in two steps: marker selection and rigorous testing before developing a final microsatellite panel for dhole individual identification.

As most number of cross-species markers was found to be from dogs and earlier used markers were less polymorphic for individual identification, we decided to first examine if both species (domestic dog and wild dog) share genetic similarity. Earlier karyotype and chromosomal banding studies [20] showed almost identical G-banding patterns, indicating high chromosomal level similarity between both species. Subsequently, we identified a total of 37 dog microsatellite loci from earlier published literature [21–26]. These markers were selected based on their polymorphism (number of alleles, PIC, observed heterozygosity etc.) and amplicon sizes in published literatures. Further, we mapped all the markers on available dog genome in UCSC Genome Browser (http://genome.ucsc.edu/; Accession ID: GCA_000002285.2) to assess the chromosome number to which each marker is associated with. Finally a total of 18 microsatellites were selected based on their amplicon size, chromosome number and polymorphism (based on published data) for further testing. The details of the markers are given in Table 1.

**Table 1:**
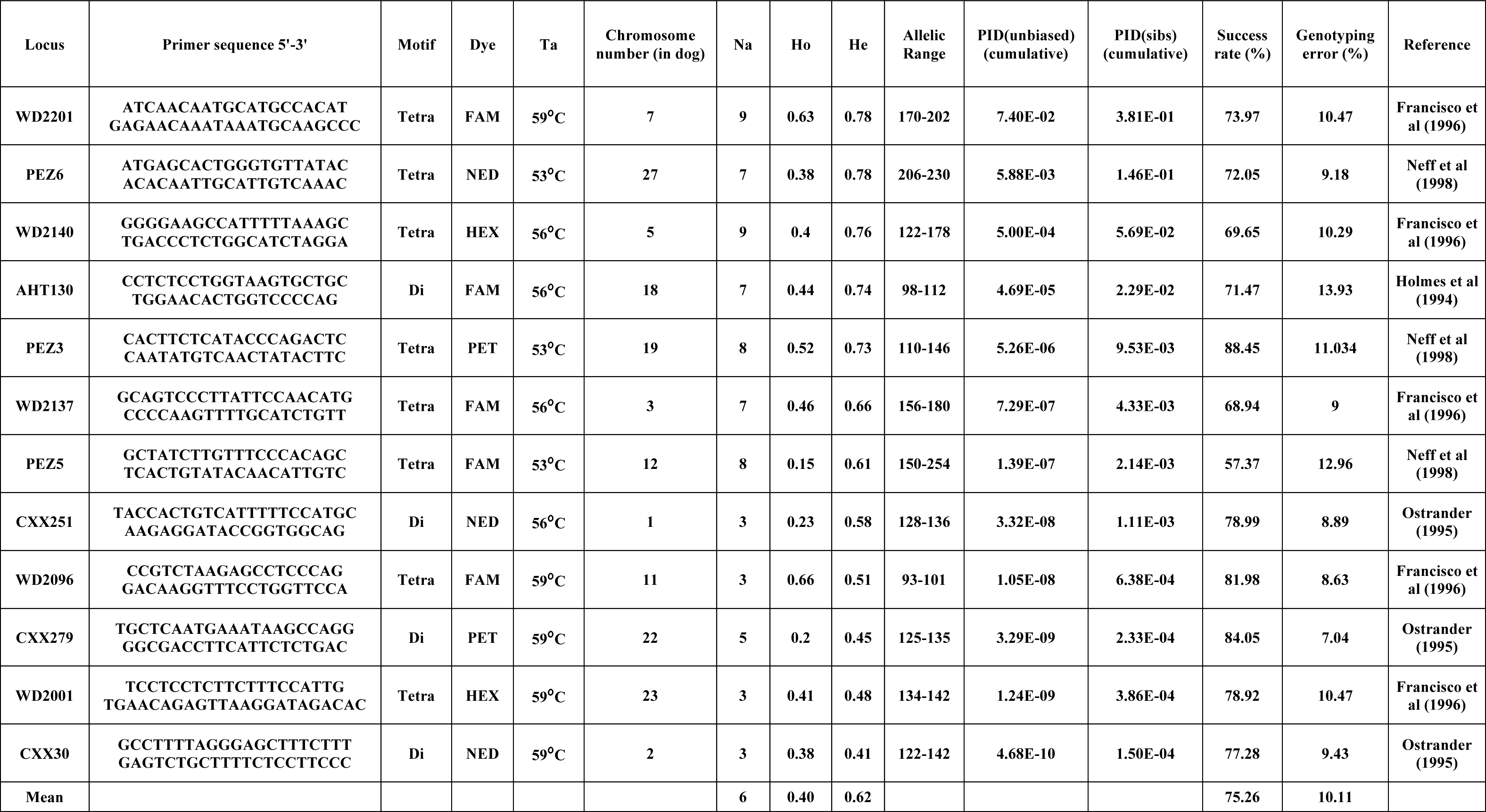
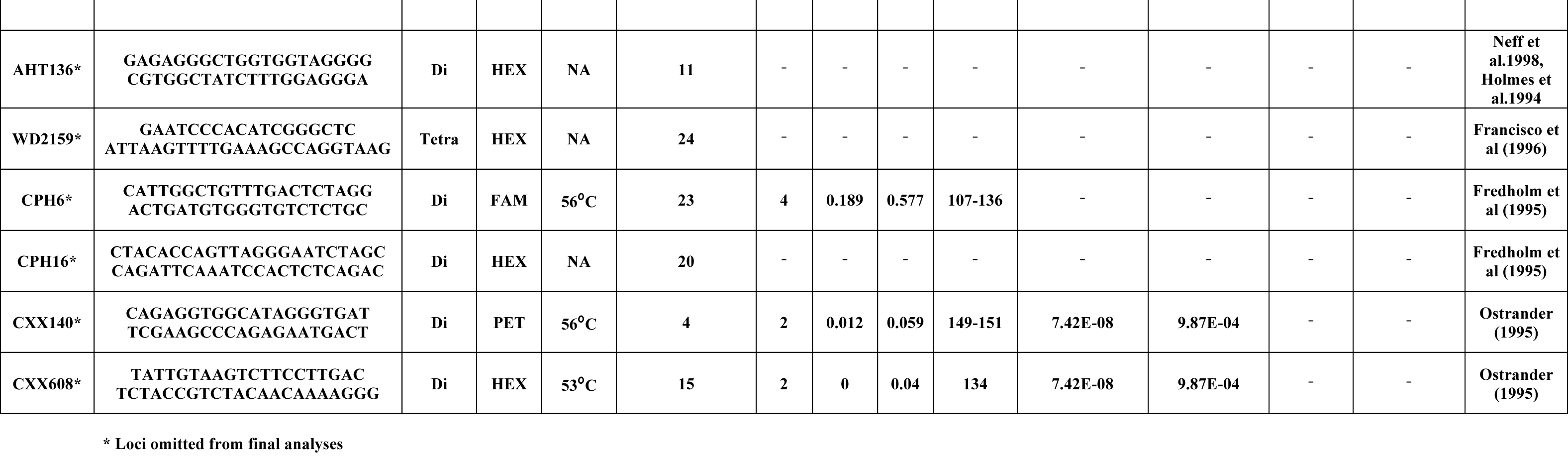
Details of microsatellite loci used in this study.

### PCR standardization and data validation

All initial standardization of the markers was conducted using dhole blood samples (n=4). PCR reactions were performed for selected 18 microsatellites in 10 μl reactions containing 3.5 μl Qiagen multiplex PCR buffer mix (QIAGEN Inc., Hilden, Germany), 0.2 μM labelled forward primer (Applied Biosystems, California, United States), 0.2 μM unlabelled reverse primer, 4 μM BSA and 2 μl of 1:50 dilution of blood DNA extract. The PCR conditions included an initial denaturation (95 °C for 15 min); 50 cycles of denaturation (94 °C for 30 s), annealing (50-60 °C gradient for 30 s) and extension (72 °C for 35 s); followed by a final extension (72 °C for 30 min). Following post-temperature standardizations markers with same annealing temperatures but with different labels or allele sizes were standardized as multiplex assays (see Table 1 for details). During all amplifications, both extraction and PCR negative controls (one PCR negative every set of 11 reactions) were included to monitor any possible contamination. Post amplification, 2 μl of PCR product was mixed with HiDi formamide (Applied Biosystems, California, United States) and LIZ 500 size standard (Applied Biosystems) and genotyped in an ABI genetic analyser (Applied Biosystems, California, United States). The fragment lengths were scored manually using the program GENEMARKER (Softgenetics Inc., Pennsylvania, Unites States). Each reaction was repeated three times to ensure good data quality.

Once the initial standardizations were performed using reference samples, final standardization was conducted with dhole faecal DNA. Species identification was performed for all field-collected faeces using the methods described in Modi et al. [27]. Further PCR reactions were conducted using the earlier standardized protocols with 1 μl of faecal DNA. Negative controls were included to monitor contaminations.

For faecal samples, data validation was performed through a modified multiple-tube approach as described in Mondol et al. [10]. All faeces that have amplified in 50% of the loci in the panel during first PCR were repeated two more times for all loci. Following allele calling, a consensus genotype was prepared using the ‘Quality index’ protocol described in Miquel et al. [28] combining all the repeats for each sample. We calculated average amplification success as the percent positive PCR for each locus, as described by (Broquet & Petit)[29]. We quantified allelic dropout and false allele rates manually as the number of dropouts or false alleles over the total number of amplifications, respectively[29], as well as using MICROCHECKER v 2.2.3 [30]. Program FreeNA [31] was used to determine the frequency of null alleles. Molecular sexing of the identified individuals was conducted using the method described in Modi et al. [27].

### Data analyses

The identity analysis module implemented in program CERVUS [32] was used to identify identical genotypes (or recaptures) by comparing data from all samples for all amplified loci. All genetic recaptures were removed from the data set. GIMLET (Valière, 2002) was used to calculate the PID_(sibs)_ for all the individuals. Following this, any allele having less than 10% frequency across all amplified loci were rechecked for allele confirmation. ARLEQUIN [34] was used to determine Hardy Weinberg equilibrium and linkage disequilibrium for all the loci.

Further, we conducted exploratory population level analyses to detect possible population structure and signatures of decline, if any, using the dhole individual genotype data generated here. For these analyses we used data from populations having information from at least 10 different individuals. We used multivariate analyses approach implemented in program Discriminant Analysis of Principal Component (DAPC) [35] to identify genetic clusters in our data. This approach transforms the genetic data into principal components, followed by clustering to define group of individuals with a consideration of minimum within-group variation and maximum between group variations among the clusters. The analyses was conducted using adegenet package 2.1.1 in R studio 1.1.453 (R Development Core Team 2018) where optimal number of clusters was determined through the Bayesian Information Criterion [35], and number of clusters was assessed using adegenet in R. Subsequently, we used a Bayesian genetic clustering approach that implemented spatial information in the form of Hidden Markov Model [36] using TESS version 1.1 [37], where the entire data was simulated for genetic subpopulations values (K) between one and ten. TESS was run for 50000 simulations (10000 burn-in) to estimate K, as well as to assign individuals to clusters.

Finally, we used two different summary statistics-based qualitative approaches to detect any signal of population decline in dholes. These approaches are the Ewens, Watterson, Cournet and Luikart method implemented in program BOTTLENECK ver 1.2.02 [38] and the Garza-Williamson index/ M-ratio approach implemented in program ARLEQUIN [34]. For BOTTLENECK, simulations were performed under three mutation models: infinite allele model (IAM), single stepwise model (SMM) and two-phase model (TPM) to obtain the distribution of H_e_ and the values are then compared to the real data values. For TPM model, 30% multi-step mutation events were allowed during the simulations. This method detects departures from mutation-drift equilibrium and neutrality, which can be explained by any departure from the null model, including selection, population growth or decline. The Garza-Williamson index uses data on the frequency and the total number of alleles and the allelic size range to investigate population decline.

## Results

During initial standardizations we tested all 18 selected markers (see Table 1) with four wild-caught dhole blood DNA samples. Three of these tested markers (WD2159, CPH16, AHT136) did not show any amplification in the blood DNA samples and were removed from subsequent analyses. The remaining 15 markers were then amplified with 225 genetically confirmed dhole faecal samples. Following data validation through multiple repeats, amplification success rates and polymorphism for these loci were calculated. The results show that loci CXX608 and CXX140 were monomorphic in all amplified samples, and locus CPH6 has low amplification success rate (~ 35%) from faecal DNA and thus were removed from the panel. The remaining 12 markers were finally standardized as four multiplex panels (see Table 1) for dhole individual identification.

None of these final 12 loci showed any signatures of large-scale allelic dropouts in MICROCHECKER. The mean allelic dropout rate was found to be 0.1. Overall frequency of null alleles was calculated as 0.037, indicating this 12 loci panel has low genotyping error rates. Overall, amplification success ranged between 57% to 88% from dhole faeces. The loci showed higher (WD2201- 9 alleles, H_o_-0.63) to medium (CXX251-3 alleles, H_o_ −0.23) levels of polymorphism (Table 1). None of the loci were found to deviate from the Hardy-Weinberg equilibrium and there were no evidences for strong linkage disequilibrium between any pair of loci. Summary statistics for various measures of polymorphism (observed and expected heterozygosity, number of alleles and allelic size range) for all loci in the final panel are presented in Table 1.

For individual identification, we only considered samples that produced good quality data for at least seven of the 12 panel loci. Out of the 225 field-collected dhole faecal samples amplified with the panel 98 produced seven or more loci data and a total of 102 samples (4 blood and 98 faecal samples) were used in downstream analyses (Figure 1). About 70% of these samples (n=71) have successfully amplified for 10-12 loci. Following analyses with CERVUS, we identified 101 unique dhole individuals from the entire data, whereas one individual from NNTR was found to be a ‘genetic recapture’. Cumulative PID_sibs_ and PID_unbiased_ values were found to be 1.5× 10^−4^ and 4.7 × 10^−10^, respectively, indicating a strong statistical support for unambiguous individual identification. The number of unique individuals from each sampled area was found to be: PTR= 35, MTR= 35, NNTR= 14, UKLWS= 7 and TATR= 10. Molecular sexing showed a success rate of 67%, with a sex ratio of 4:1 in all identified dhole individuals (n=101).

**Figure 1:**
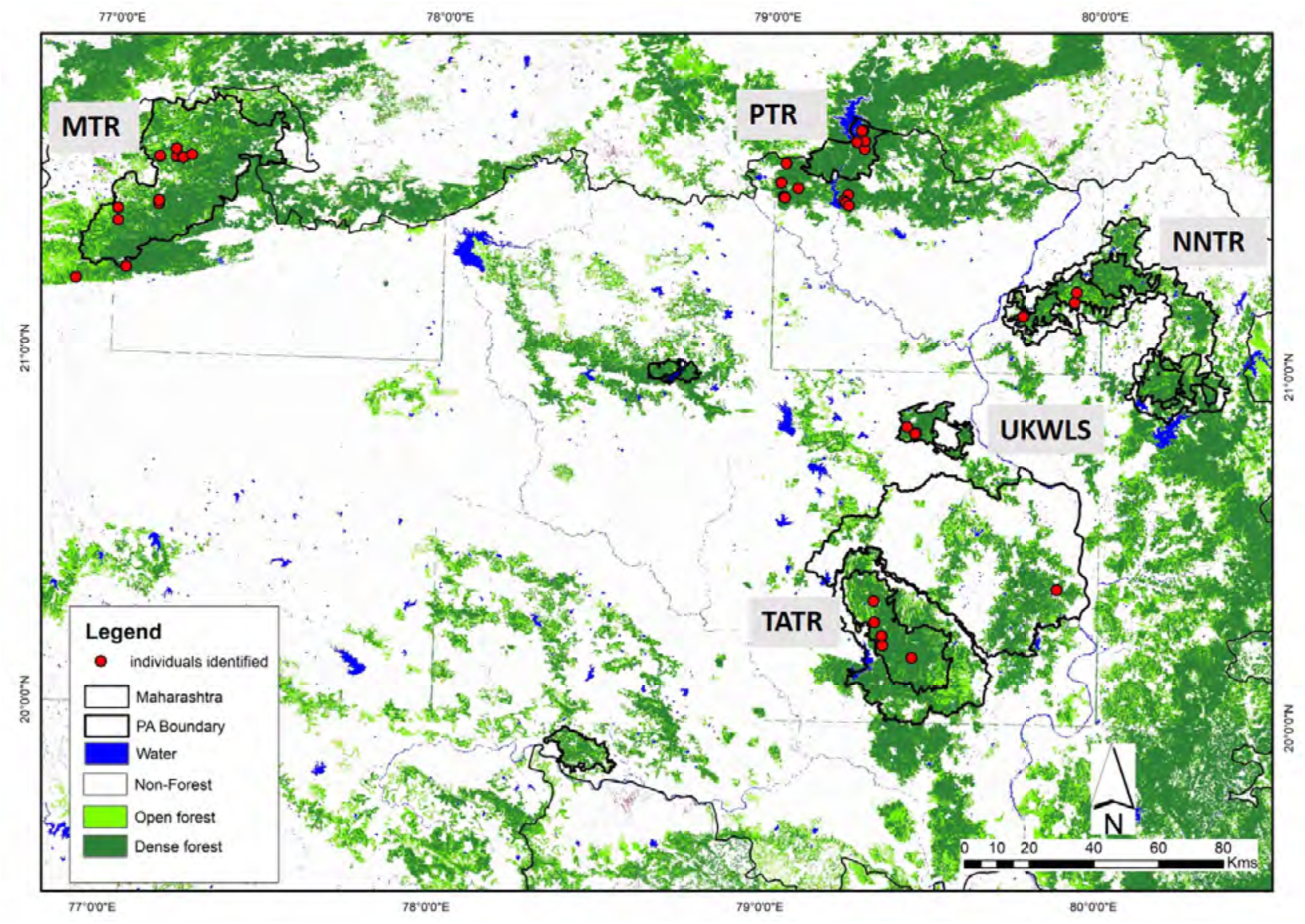
Map of the protected areas in the study with locations of number of individuals identified.

Our exploratory analyses with DAPC identified six genetic clusters within the individual dholes from all sampled areas (n=101). The first principal axis indicated the separation between PTR and MTR-NNTR and the second principal axis separated MTR and NNTR (Figure 2a). The clusters indicate no genetic sharing between MTR and PTR, and NNTR individuals are genetically closer to both MTR and PTR. Similarly, the Bayesian clustering approach identified four genetic clusters of MTR, PTR, TATR and NNTR with mixed genetic ancestry in UKWLS (Figure 2b).

**Figure 2a:**
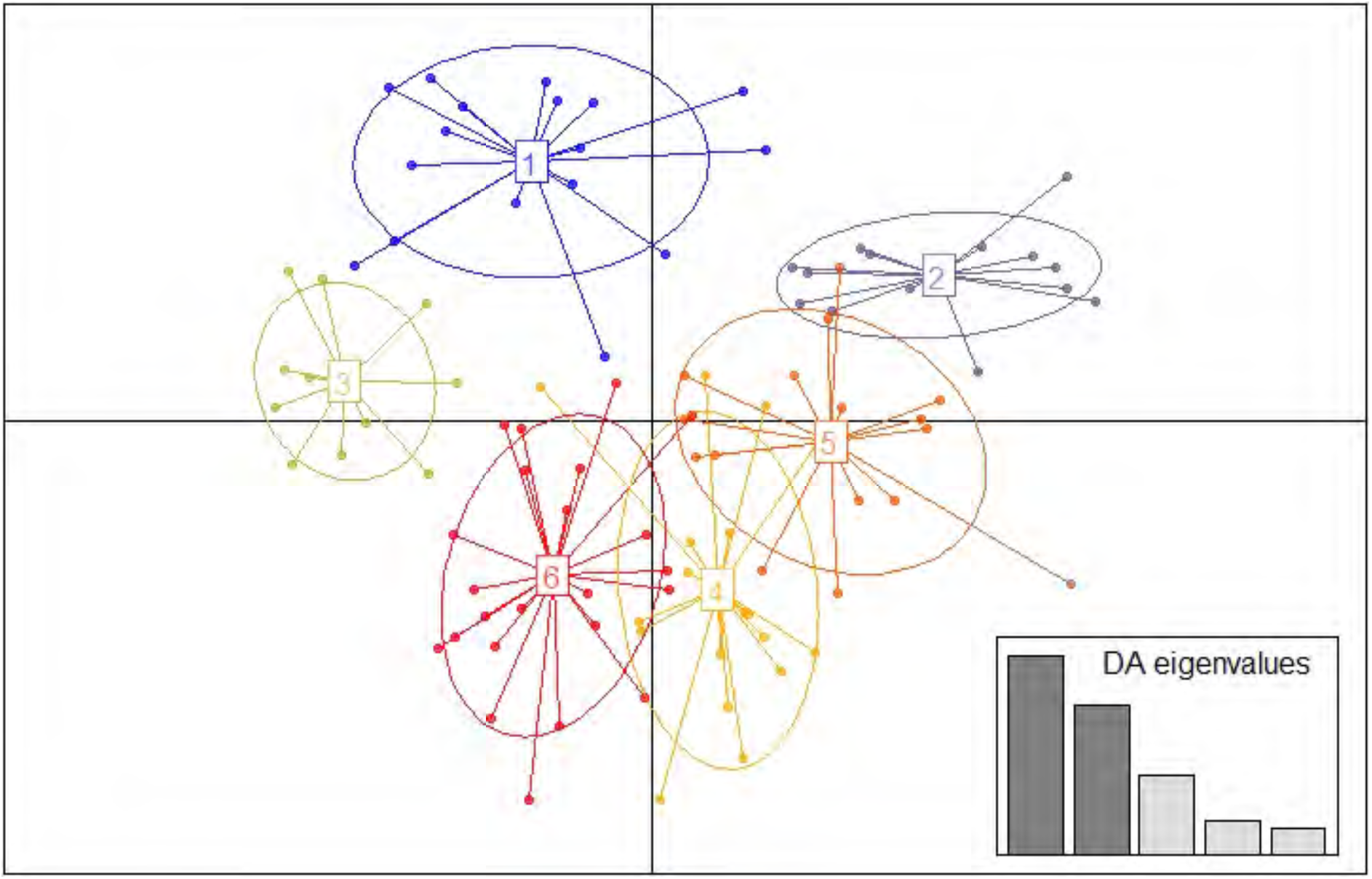
Discriminant Analysis of Principal Components (DAPC) of microsatellite markers for 102 individuals of *Cuon alpinus*.

**Figure 2b:**
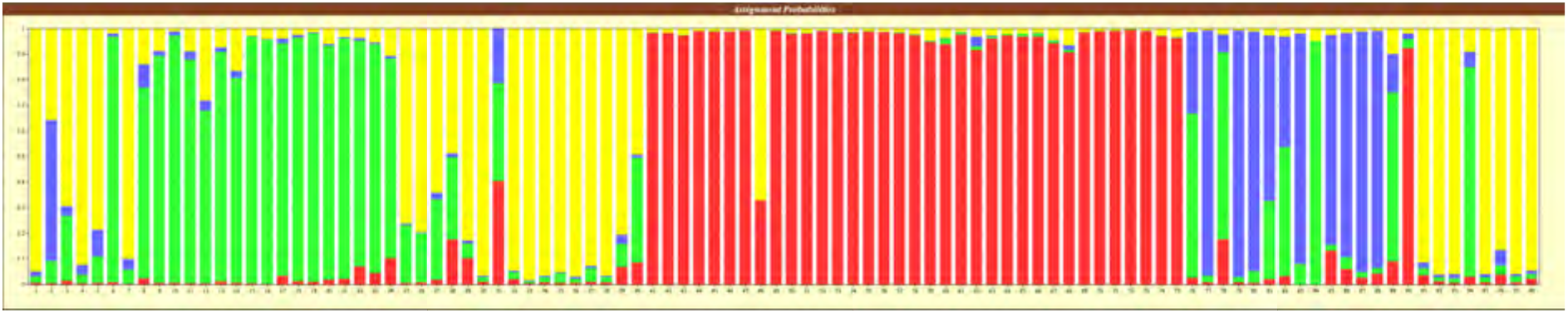
Results of genetic structure of dhole individuals from four protected areas in Maharashtra, India.

Population decline analyses in dholes from PTR and MTR detected heterozygote excess in 11 of the 12 loci depending on the mutation models used, suggesting a loss of rare alleles through population decline for all three populations. Similarly, the M-ratio approach shows a low ratio between number of alleles (NA) and the allelic size range in all populations (M-ratio PTR- 0.27611 (SD 0.12); M-ratio MTR- 0.35809 (SD 0.24)), indicating signals of population decline.

## Discussion

In this paper, we standardized protocols for individual identification of Asiatic wild dogs from poor quality DNA samples, and the final marker panel could unambiguously identify individual dholes in our field-based pilot study from five protected areas of Maharashtra, India. The systematic protocols followed here offer some advantages over earlier efforts on dhole individual identification from faecal samples by Iyengar et al. [19]. Firstly, use of a large panel of 37 microsatellite loci for preliminary assessment of marker suitability along with genomic mapping-based selection of final markers (n=18) allowed us to ascertain a combination of loci for unambiguous individual identification with high statistical power. The rigorous testing of the loci with large number of DNA samples from different sources also allowed us to exclude loci that might be problematic due to low amplification success from non-invasive samples. The final panel consisting 12 markers were further standardized into four multiplex reactions to provide time and cost-effective options during data generation. We were very careful to initially select a large number of tetranucleotide markers as they are known to have low stutter peak problems and better allele characteristics from poor quality samples [39], while dinucleotide markers generally have higher amplification success [40]. Thus, our final panel with a ratio of 2:1 tetra vs. dinucleotide microsatellites would provide the ideal combination in terms of high success rate and less technical issues in allele calling during dhole individual identification. The amplification success rate for all loci was > 70% except locus PEZ5 (~ 60%), but it was found to be polymorphic and was included in the panel. The overall genotyping error frequency was found to be < 0.2 from dhole faeces, which is within the recommended limits for non-invasive population genetic research [41].

Our motivation in this study was to develop effective protocols that could be applied for individual identification of Asiatic wild dogs as they are difficult to identify from physical characteristics (spots, marks, stripes etc.). Their elusive nature also makes it challenging to estimate population size using traditional techniques (photographic capture, field-based observations etc.) at landscape levels. For genetic estimation of population size Waits et al. (2001) recommended a threshold PID_sibs_ value that is at least double than the approximate number of animals in any given area. Given the global dhole population estimate of 949-2,215 mature individuals [4] and the cumulative PID_sibs_ value of 1.5× 10^−4^ achieved in this study, this panel could be used for dhole population estimation at a landscape level. However, it is important to point out that we generated individual level information from about 43.5% (98 out of 225 faeces) of the field-collected samples in this study. Similar patterns of low amplification success rate from field-collected faecal samples have been observed in earlier genetic study of dhole [19], leopard [42] and other species [40]. Considering dhole cryptic nature, social behaviour and ecology in corroboration with low amplification success rate, we suggest an intensive faecal sampling effort for estimation of population size for this species. We have identified only one genetic recapture from the field-collected faecal samples, but this could be attributed to our sampling strategy to cover large geographical area and maximize collection of faeces from potentially different individuals.

Although our sampling strategy was not exhaustive enough to assess detailed population genetic parameters for dholes, our exploratory analyses showed some interesting patterns with the individual level data. Both DAPC and TESS results indicate no genetic connectivity between dhole populations of MTR and PTR, while NNTR individuals show genetic similarity both populations. These patterns are also described in earlier studies where data from multiple species indicated little gene flow between PTR and MTR [43–46] and connectivity between NNTR and MTR [47]. The most likely reason behind such pattern in dholes could be severe defragmentation and loss of forest patches in recent times, and dhole being obligate forest-living species possibly cannot disperse between these two protected areas. However, further extensive sampling covering PTR, MTR, NNTR and Satpura-Maikal landscape can confirm this pattern. Similarly, two separate qualitative analyses of dhole demography indicated signals of population decline in both PTR and MTR. Further quantitative analyses after landscape level sampling will help us to assess the extent and timing of such decline [48, 49]

## Conclusion

In the broader context of understanding dhole population dynamics at local or landscape scales, genetic sampling is possibly the only way to generate information with spatial and temporal coverage for this elusive, social carnivore as photographic sampling or conventional tagging cannot be employed due to lack of distinguishing natural marks and logistical difficulties of physical captures of large number of animals. Results from this study provide a great tool to generate individual level information from field-collected faecal samples. In combination with a good sampling strategy, our methods can be used in a cost-effective way to investigate species biology (including patterns of genetic diversity, relatedness and population connectivity) as well as to estimate population abundance of dholes in the wild.

## Acknowledgement

We are thankful to the Maharashtra State Forest Department for providing research permission, and the forest officials and staff for their assistance in fieldwork. We thank the Dean, Faculty of Wildlife Sciences and Director, WII for their continuous support in this research. We thank A. Madhanraj, Tanima and Aditi for their help in the laboratory and Shaheer for his help with GIS. This research was funded by Maharashtra State Forest Department and National Tiger Conservation Authority, Government of India. S. Modi was supported by the CSIR SRF fellowship and SM was supported by the INSPIRE Faculty Award (LSBM-47) of Department of Science and Technology, Government of India.

## Author’s contributions

BH and Samrat Mondol have conceived the idea. Shrushti Modi, PG and PN collected the samples and associated data. Shrushti Modi conducted the laboratory experiments. Shrushti Modi and Samrat Mondol conducted the analyses. All authors commented on the final manuscript.

---

**Supplementary Figure 1:**
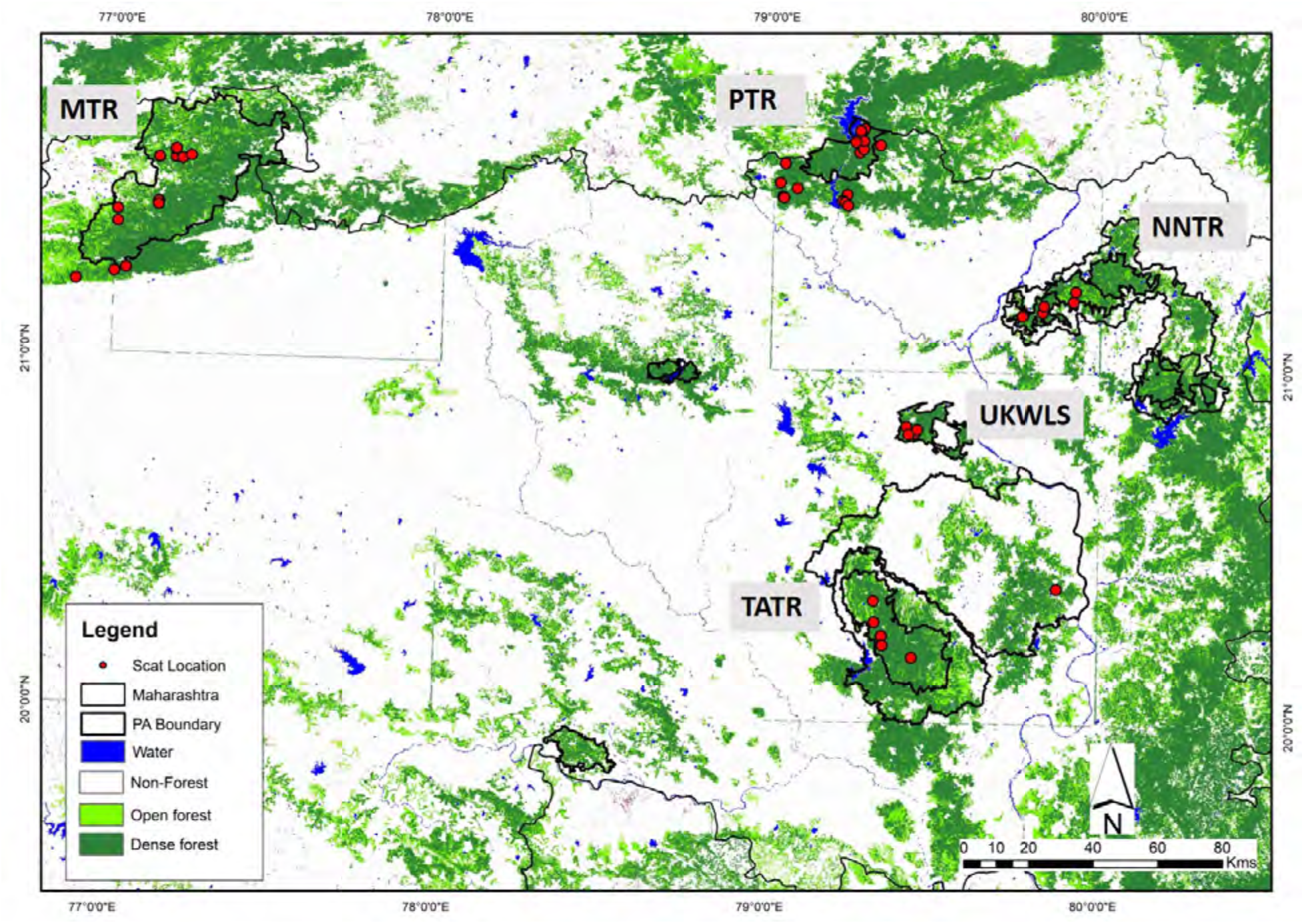
Map of the protected areas in the study with locations of scat samples confirmed as dholes.

